# Mitochondrial dysfunction reshapes methyl-group allocation in skeletal muscle

**DOI:** 10.64898/2026.08.23.746543

**Authors:** Anastasiia Marmyleva, Ville Tiusanen, Priit Joers, Biswajyoti Sahu, Anu Suomalainen

## Abstract

Mitochondria are central metabolic organelles with functions extending beyond energy production to anabolic and folate-mediated one-carbon (1C) metabolism. One-carbon metabolism supports methylation reactions that modify diverse targets including metabolites, nucleic acids, and chromatin, and has emerged as a contributor to mitochondrial disease-related stress responses. Here, we report tissue-specific remodeling of methylation events in response to mtDNA replication defect, using the deletor mice carrying a dominant mutation in Twinkle, the replicative helicase of mtDNA, causing adult-onset mitochondrial myopathy (MM) in humans and mice. In affected skeletal muscle, deletors show a distinct methylation signature, with increased creatine synthesis and reduced phosphatidylcholine production, two major consumers of S-adenosylmethionine-derived methyl groups. We further observed tissue-specific upregulation of selected RNA methylation marks and redistribution of the repressive histone mark H3K9me3, indicating coordinated remodeling of metabolic and epigenetic methylation pathways. Our evidence shows that a mtDNA replication defect remodels muscle-specific methylation signature of phospholipids, histones and RNA, identifying methylation remodeling as a key component of MM pathogenesis.

## Introduction

Recent years have witnessed a switch in the common perception of mitochondria as simple energy-generating organelles with a growing recognition of their role as central metabolic hubs. Diseases that originate from such mitochondrial malfunctions are highly variable, sometimes affecting multiple organs, and sometimes restricted to a specific tissue. The diseases trigger molecular metabolic responses both locally and systemically, the types of which depend on the original molecular defect and the affected tissue (Suomalainen & Battersby, 2018). How the remodeled metabolic pathways contribute to the high variability of clinical manifestations of mitochondrial diseases is a big open question in the field (reviewed in Suomalainen & Nunnari, 2024).

Mitochondrial myopathy (MM) is an adult-onset disease that can be caused by several different genetic defects either in the nuclear genome or mitochondrial DNA (mtDNA). The typical causes include mtDNA deletions, either single or multiple, mitochondrial tRNA mutations, all affecting mtDNA expression and protein synthesis. What determines such sensitivity of muscle to such defects is unclear. Previous studies have characterized a model for MM, the deletor mouse, carrying a dominant patient mutation in mtDNA replicative helicase Twinkle that results in slowly accumulating multiple mtDNA deletions and progressive deficiency of the mitochondrial respiratory chain in the skeletal muscle, heart and specific regions of the hippocampus of the brain (Tyynismaa et al, 2005). These mice as well as the patients with the same mutation present a wide-spread transcriptomic and metabolic response, together called mitochondrial integrated stress response (ISRmt) (Tyynismaa et al, 2010; Nikkanen et al, 2016; Khan et al, 2017; Forsström et al, 2019; Pirinen et al, 2020). The metabolic component of ISRmt remodels the one-carbon (1C) metabolism, the major biosynthetic pathways providing nucleotides, phospholipids, redox components, amino acids and methyl groups for cellular methylation reactions in a cell-type specific manner (Nikkanen et al, 2016; Bao et al, 2016; Kühl et al, 2017). The deletor studies have indicated that in MM the glucose-derived 1C atoms are directed to glutathione synthesis and increase purine synthesis (Nikkanen et al, 2016). However, the role of methyl cycle, with its cycled conversion between the universal methyl donor – S-adenosylmethionine (SAM) - into its byproduct – S-adenosylhomocysteine (SAH) – has remained unknown.

Transformation of SAM into SAH is accompanied by the release of a methyl group and its donation to many kinds of acceptors that are also called “sinks”: from other metabolites to proteins like histones, to nucleic acids (DNA and RNA) (Mudd et al, 2007; Karimian et al, 2023). The methyl-group sinks in the skeletal muscle are synthesis of creatine involved in energy metabolism; phosphatidylcholine, an essential membrane lipid; sarcosine, a methylated intermediate of glycine metabolism, and methylated arginine derivatives, including symmetric and asymmetric dimethylarginines (SDMA and ADMA) released during the turnover of arginine-methylated proteins (Mudd et al, 2007; Schlesinger et al, 2016).

Here, we report that mitochondrial myopathy induces methyl cycle rearrangement. The deficient lipid biosynthesis, particularly, of phosphatidylcholine, as well as creatine leave a pool of free methyl groups available for new donors. We show accumulation of repressive trimethylation mark at lysine 9 of histone H3 (H3K9me3) and specific posttranscriptional modifications of RNA. These changes were not present in the liver of the same mice, highlighting the tissue-specific nature of methyl cycle remodeling in MM.

## Results and discussion

### Limited methyl group supply to phosphatidylcholine synthesis in mitochondrial myopathy

To evaluate changes in the metabolic part of the methyl cycle in response to mitochondrial muscle disease, we analyzed the RNA expression profile of the enzymes that catalyze these reactions and a targeted metabolomics analysis of deletor skeletal muscle. The muscle presented a decrease in expression of phosphatidylethanolamine N-methyltransferase (PEMT) and phosphoethanolamine (PEA), a precursor of PEMT substrate phosphatidylethanolamine **(Fig. 1A-C)**. The elevated PE (phosphatidylethanolamine) to PC (phosphatidylcholine) ratio suggests a decline in PC synthesis **(Fig. 1D)**. Further, the mice showed a near 50% decrease in guanidinoacetate N-methyltransferase (GAMT) required to methylate guanidinoacetate (GAA) to creatine using SAM as a methyl donor **(Fig. 1A,E)**. The low GAMT expression, combined with increased creatine synthesis precursor GAA, increased creatinine and the trend of decreased creatine to creatinine ratio all point to decreased availability of methyl groups for creatine synthesis, despite the 1.3-fold elevated steady-state creatine level **(Fig. 1F)**. Similar changes in phospholipids or creatine were not observed in liver tissue **(Fig. 1G, Fig. EV1A)**.

**Figure 1.**
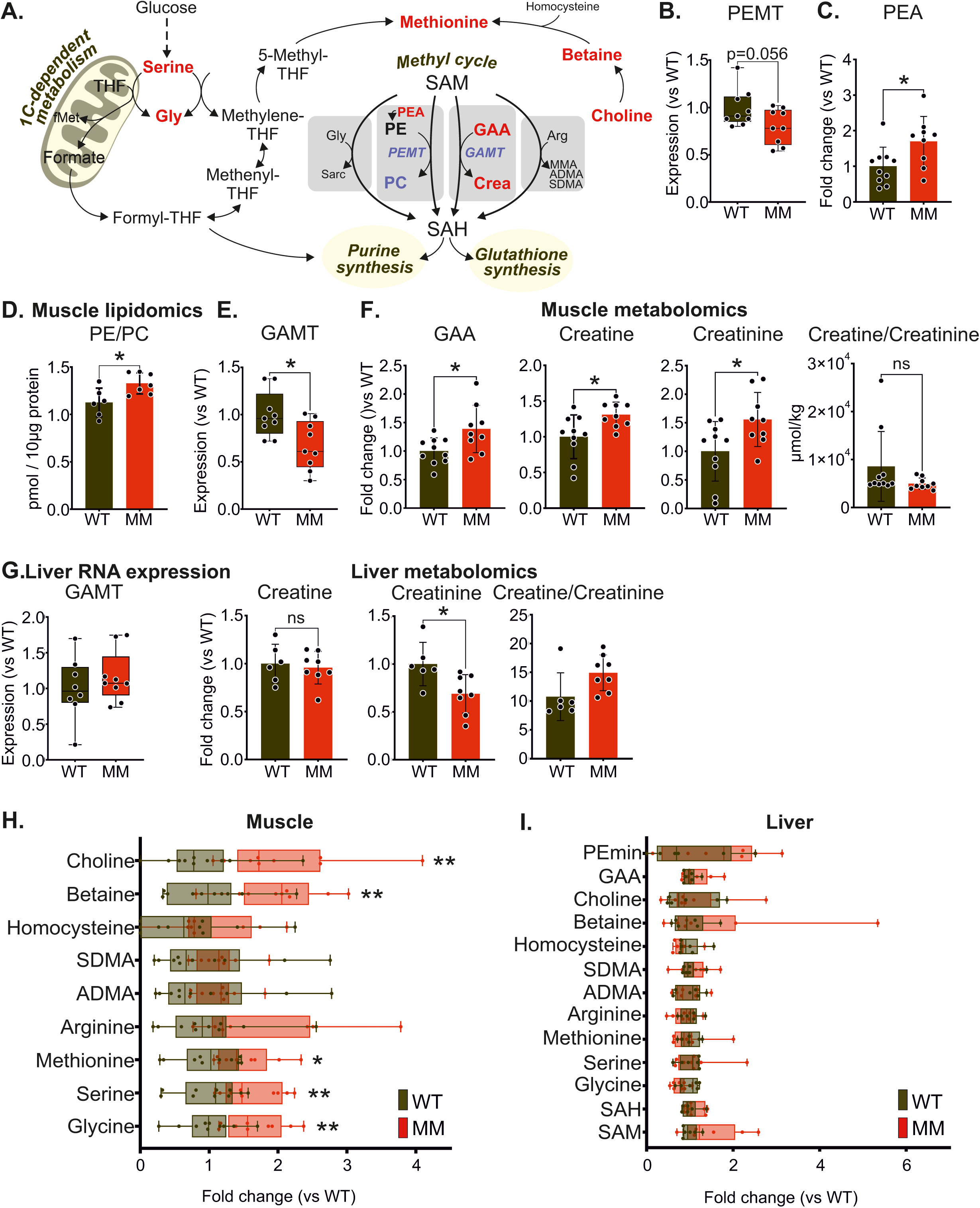
– Comparing metabolic rearrangements of methyl groups between skeletal muscle and liver of MM mice. **(A)** Mitochondrial and cytoplasmic branches of one carbon metabolism and individual metabolic methylation reactions. Blue, decreased amount; red – increased amounts of metabolites or enzyme-encoding genes. **(B)** mRNA expression of PEMT enzyme in skeletal muscle (n = 9 mice/group). **(C)** Metabolic levels of PEA in skeletal muscle; liquid chromatography and mass spectrometry (n = 9 mice/group). **(D)** Ratio between PE and PC phospholipids in skeletal muscle; lipidomics assay (n = 6-7/group). **(E)** mRNA expression of GAMT enzyme in skeletal muscle (n = 9 mice/group). **(F)** Levels and ratio of creatine metabolic intermediates in skeletal muscle; liquid chromatography and mass spectrometry (n = 9 mice/group). **(G)** mRNA expression of GAMT enzyme and metabolic levels and ration of creatine and creatinine in liver (n = 6-9/group). **(H – I)** Levels of methyl-cycle-related metabolites in **(H)** skeletal muscle and **(I)** liver (n = 9/group). Abbreviations: Gly, glycine; THF, tetrahydrofolate; fMet, formylmethionine; SAM, S-adenosylmethionine; SAH, S-adenosylhomocysteine; GAA, guanidinoacetic acid; Crea, creatine; PEA, phosphoethanolamine; PE, phosphatidylethanolamine; PC, phosphatidylcholine; GAMT, guanidinoacetate N-methyltransferase; PEMT, Phosphatidylethanolamine N-methyltransferase; PEmin, phosphoethanolamine; SDMA, symmetric dimethylarginine; ADMA, asymmetric dimethylarginine; WT, wild type; MM, mitochondrial myopathy. Data information: Gene expression data and metabolites in F,D are presented as 25^th^ and 75^th^ percentiles with whiskers spanning to min and max points. Other metabolite scatter plots visualize mean with standard deviation (SD) *P≤0.05, **P≤0.01, ***P≤0.001, ****P≤0.0001 (Student’s t-test)

Creatine synthesis is commonly understood as an organ-compartmentalized process: the kidney produces the precursor GAA, which the liver then converts into creatine. Afterwards, newly produced creatine is supplied to high-demand tissues such as skeletal muscle. In our model, however, creatine production is supported by muscle itself. Hepatic GAMT expression remained stable, which indicates that the liver did not adjust its creatine supply in response to muscle disease **(Fig. 1G)**.

Methyl donor metabolites that support methionine regeneration and methyl cycle activity, including choline, betaine, serine, as well as methionine itself, increased in the muscle of deletors but not in their liver **(Fig. 1H,I)**. Liver, however, showed a lowered adenosylhomocysteinase (AHCY) expression **(Fig. EV1A)** which suggests accumulation of SAH that prevents methylation reactions. The methionineadenosyltransferase-2-alpha (MAT2a) and AHCY, related to production of sarcosine and modified versions of arginine respectively, remained stable in muscle **(Fig. EV1B)**. Similarly, liver-specific MAT1A also did not show any changes **(Fig. EV1A)**. The metabolic products of arginine (ADMA or SDMA) nor sarcosine metabolism were not covered in our dataset.

Altogether, the evidence suggests tissue-specifically modified methylation reactions in mitochondrial myopathy. Muscle-specific findings were also preserved in human MM patients, who also showed increased metabolic components and precursors of the methyl cycle (Buzkova et al, 2018). Also, methionine was one of the most significantly increased targets in muscle and blood of MM patients, and the blood showed high levels of betaine. Increased GAA in the MM blood supports decreased creatine synthesis (Buzkova et al, 2018).

### Histone 3 trimethylation (H3K9me3) becomes a major methyl sink in MM muscle

Prompted by the decreased methyl sinks in the deletor muscle, we asked whether methyl groups remained available for chromatin, specifically the repressive H3K9me3 mark. Previous study in human cancer cells reported H3K9me3 as a repressor of ISR components such as ATF4 and genes for serine (PHGDH and PSAT1) and glutathione (CTH) synthesis among others (Zhao et al, 2016). The close resemblance of ISRmt in mitochondrial myopathy to the cancer cells response motivated us to test the repressor mark in muscle disease.

Total chromatin extracts revealed mild increase of H3K9me3 immunopositivity in MM muscle compared to healthy tissue, a proportion evaluated against total histone H3 **(Fig. 2A, B)**. This stimulated us to closely assess the genomic distribution of this mark with ChIP-sequencing method. PCA analysis of sequencing results showed a clear separation between control (WT) and disease (DEL) groups **(Fig. 2C)** and the number of H3K9me3-specific peaks was higher in MM muscles compared to the controls **(Fig. 2D)**. The results indicate that chromatin becomes a preferred methyl group sink in the skeletal muscle with mitochondrial disease.

**Figure 2.**
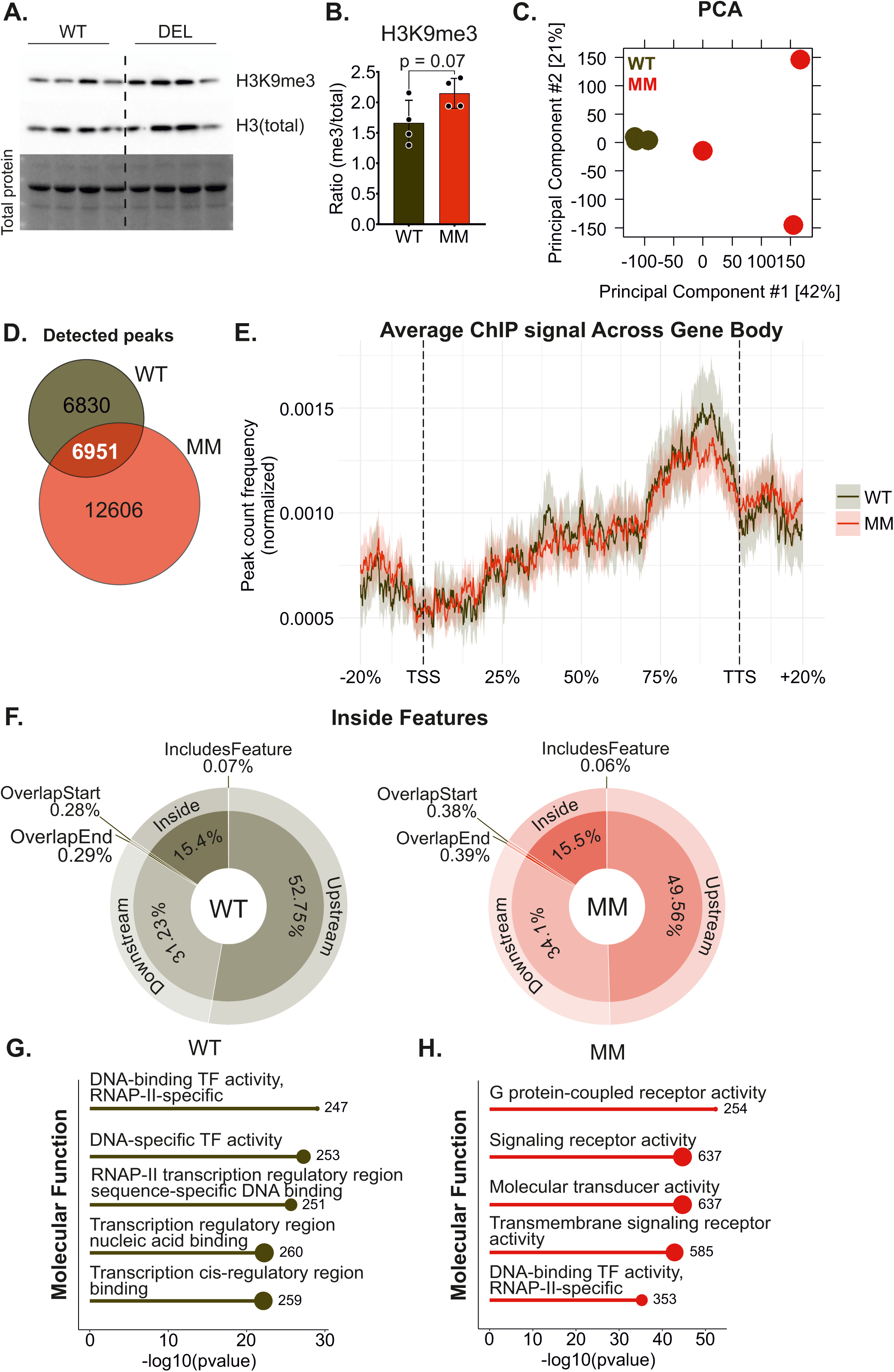
– Distribution of repressive H3KGme3 chromatin mark across skeletal muscle genome in MM mice. **(A)** Protein levels of methylated histone H3 - H3K9me3 - and total H3 in skeletal muscle detected by immunoblotting. **(B)** Ratio between H3K9me3 and total H3, quantification of (A). **(C)** PCA analysis for ChIP-seq mouse muscle samples. **(D)** Venn diagram representing unique and shared H3K9me3 modified peaks detected in ChIP-Seq assay. **(E)** Distribution of H3K9me3 signal across gene body; TSS, transcription start site; TTS, transcription termination site. **(F)** Distribution of H3K9me3 ChIP signaling across genomic features. **(G – H)** Gene ontology analysis for Molecular Function of detected ChIP-Seq peaks. Abbreviations: WT, wild type; MM, mitochondrial myopathy. n = 3/group.

### H3K9me3 enrichment in different genomic regions is modified under muscle disease conditions

H3K9me3 is well-known for being a mark of constitutive heterochromatin as it compacts DNA, thus making it inaccessible to transcriptional factors (Lachner et al, 2001; Peters et al, 2003; Verschure et al, 2005). It particularly accumulates at repetitive DNA regions such as pericentromeres, telomers, transposons and such (Tatarakis et al, 2025). Within the genome structure, H3K9me3 predominantly localizes outside of gene-coding regions and to a smaller extent withing gene body (Lee et al, 2020; Hada et al, 2022).

The average profile of H3K9me3 signal distribution between Transcription Start Site (TSS) and Transcription Termination Sites (TTS) confirmed the general pattern of this mark: its binding gradually increased closer to TTS with the maximum at the 75 percentile and the peak being smaller in MM **(Fig. 2E)**. Looking at bulk peaks detected for both MM and WT muscles, we observed that only 15% of this mark localized inside the gene coding regions whilst the strongest preference was for upstream and downstream non-coding elements **(Fig. 2F)**. The extent of such preferences varied depending on the genotype with slightly more affinity to upstream regions in WT and downstream regions in MM tissues **(Fig. 2F)**.

Even such slight redistribution of H3K9me3 peaks was sufficient to reflect the differences in the associated molecular functions as evaluated by gene ontology (GO) terms. In healthy muscles, H3K9me3 showed the strongest enrichment for regions related to transcription and transcription factor binding **(Fig. 2G)**. On the other hand, while myopathy also presented an affinity towards transcription-related genes, it enhanced correlation of this mark with signaling and receptor activity **(Fig. 2H)**. GO terms for Biological Process and Cellular Component remained the same between the groups with almost exclusive association with synapses (organization, assembly and function; **Fig. EV2A,B**).

These findings indicate that muscle increases deposition of methyl groups in myopathy particularly within the distal genomic regions with a majority of genes with modified H3K9me3 abundance belonging to G protein coupled receptors of synapses. This prompted us to further evaluate if these changes carry functional implications and result in altered expression of the relevant genes.

### H3K9me3 accumulates in genes related to synapse formation

To identify unique H3K9me3 regions that are significantly altered in myopathy compared to healthy muscle, we analyzed the ChIP-seq data using DiffBind (Stark & Brown, 2011). As expected, MM muscles showed a higher number of regions with increased H3K9me3 signal (UP, red) compared to those with decreased signal (DOWN, blue) **(Fig. 3A,B)**. A similar analysis of differentially bound peaks revealed that these changes predominantly occur in distal intergenic and intronic regions of genes **(Fig. 3C)**.

**Figure 3.**
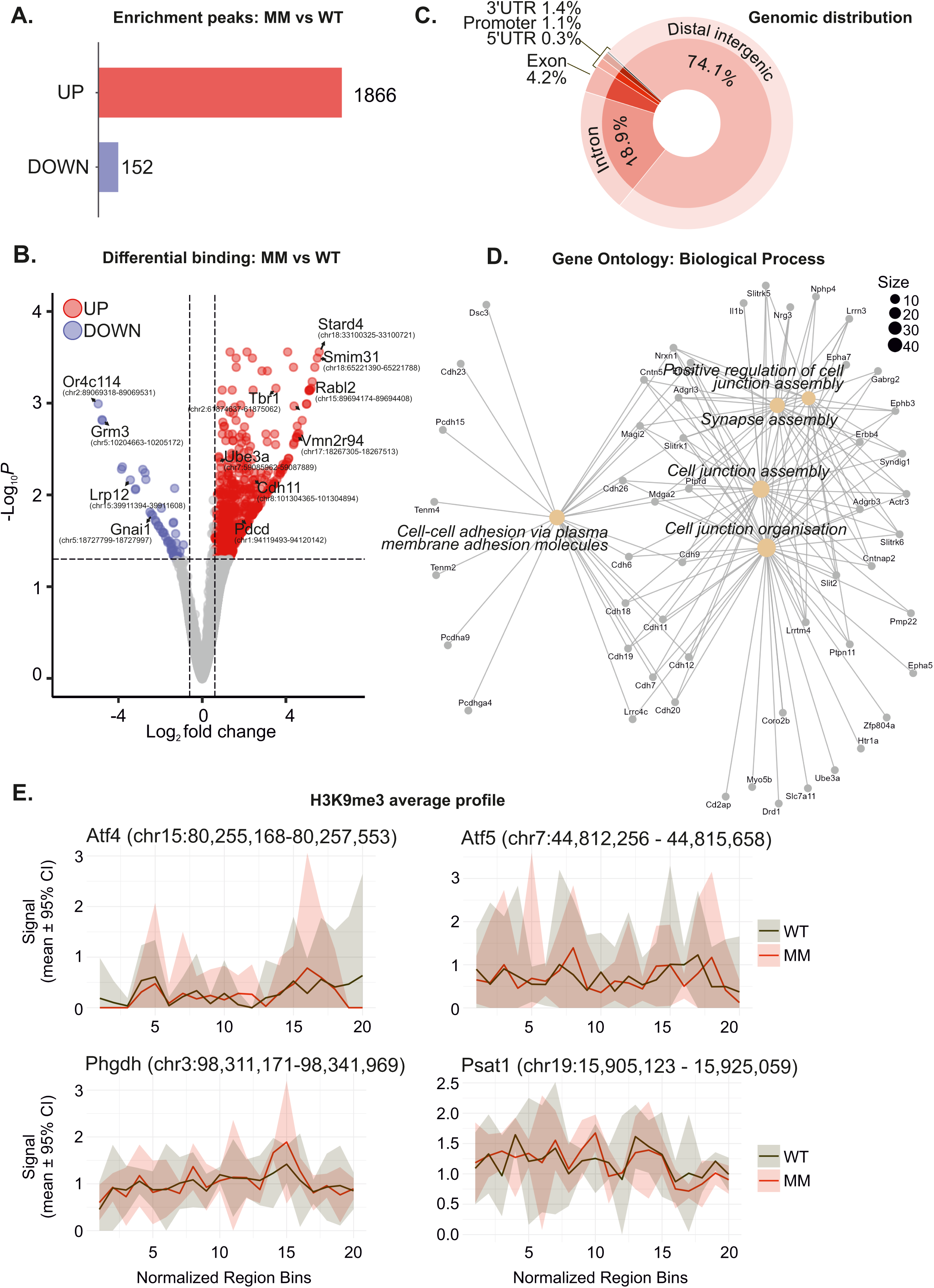
– Differential rearrangements of H3KGme3 in muscle under mitochondrial replication stress. **(A)** Differential H3K9me3 peaks showing regions with decreased (DOWN) or increased (UP) enrichment in MM muscles compared to WT. **(B)** Volcano plot of differentially bound H3K9me3 regions in MM muscles. **(C)** Genomic distribution of differentially bound regions across gene-associated features. **(D)** Gene ontology enrichment analysis differentially modified regions. **(E)** Metagene profile of H3K9me3 occupancy across gene bodies. Data represented as mean signal with 95% confidence interval. Abbreviations: WT, wild type; MM, mitochondrial myopathy. n = 3/group.

Functionally and consistent with our analysis of total peaks, GO enrichment of differentially modified regions also associated with synapse-related terms and interconnected signaling pathways **(Fig. 3D; EV2C)**. Considering the involvement of ISRmt-related processes and the known role of H3K9me3 in regulating ISRmt gene expression, we expected a reduction in H3K9me3 binding in our dataset. However, disease status did not significantly alter H3K9me3 binding at ATF4, ATF5, PHGDH and PSAT1 **(Fig. 3E)**, which are primary targets of this modification under mitochondrial stress in human 2D cultures (Zhao et al, 2016).

Collectively, these results indicate that in mitochondrial myopathy H3K9me3 is redistributed, accumulating predominantly in non-coding regions associated with synapse-related group of genes and does not appear to be required for ISRmt gene expression. Chromatin-state annotation of the up- and downregulated peaks revealed that over 70% of both sets mapped to heterochromatin, consistent with H3K9me3 being a canonical heterochromatin marker (reviewed in Bannister & Kouzarides, 2011). Upregulated regions were, to a lesser extent, also distributed across assembly gaps and artifacts (mGapArtf) with H3K9me3 association, quiescent chromatin (mQuies), zinc finger genes (mZNF), and weak enhancers (mEnhWk) (**Fig. EV2D**). Among downregulated peaks, mQuies and mGapArtf were the next most represented chromatin states (**Fig. EV2D**).

Previous studies have established a role of H3K9me3 in the control of cell identity. In human fibroblasts, H3K9me3-marked heterochromatin acts as a barrier to reprogramming by limiting binding of Yamanaka factors to specific genomic regions (Soufi et al, 2012). Similarly, in *C.elegans*, H3K9 methylation contributes to transcriptional control by restricting the availability of genomic regions to transcription factors, thereby maintaining differentiated cell identity. In addition, H3K9me3 and H3K9me2 exhibit partially redundant effects on gene repression in differentiated cells, where loss of H3K9me3 alone is not sufficient to derepress silenced genes, suggesting that demethylation can maintain gene silencing in its absence. These findings further suggests that H3K9me3 is typically deposited after H3K9me2, acting as a secondary reinforcing layer of transcriptional repression (Methot et al, 2021). In our model, H3K9me3 may similarly serve as a reinforcing mark while also acting as a methyl group sink that diverts methyl groups away from competing methyl-consuming reactions.

These findings contrast with in vitro data in human neuroblastoma BE(2)-C cells, where loss of H3K9me3 at the ATF4 locus increases its transcription. Newly produced ATF4 protein further recruits H3K9me3 demethylases to de novo serine synthesis genes, promoting their transcriptional activation (Zhao et al, 2016). This discrepancy could be partially explained by context-dependent regulation of ISRmt. Although ATF4 is upregulated in MM muscles, ATF5 is the main factor driving ISRmt response in the terminally differentiated muscle cells, whereas ATF4 plays the principal regulatory role under mitochondrial stress in cultured cells (Forsström et al, 2019; Jackson et al, 2025). Thus, the effects of H3K9me3 on ISRmt gene expression may be more pronounced in proliferating cells.

### MM selectively increases methylation of certain RNA modifications in muscle but not in liver tissues

Since methylation reactions modify nucleic acids, RNA molecules can also appear as targets of these modifications. We profiled total RNA from skeletal muscle and liver for a complex of different methylation modifications. Since we did not perform separation of types of RNAs, our fractions were predominantly enriched in rRNAs and tRNAs.

We detected a small but significant increase in two RNA marks in MM: 2’-O-methylguanosine (Gm) and a peak consistent with C2-methyladenosine (m2A) **(Fig. 4A, B)**. This increase was statistically significant in the tissue severely affected by the disease – skeletal muscle – whereas the unaffected liver showed no change **(Fig. 4C)**. Other RNA methylation sites were unaffected in both organs **(Fig. EV3 A,B)**.

**Figure 4.**
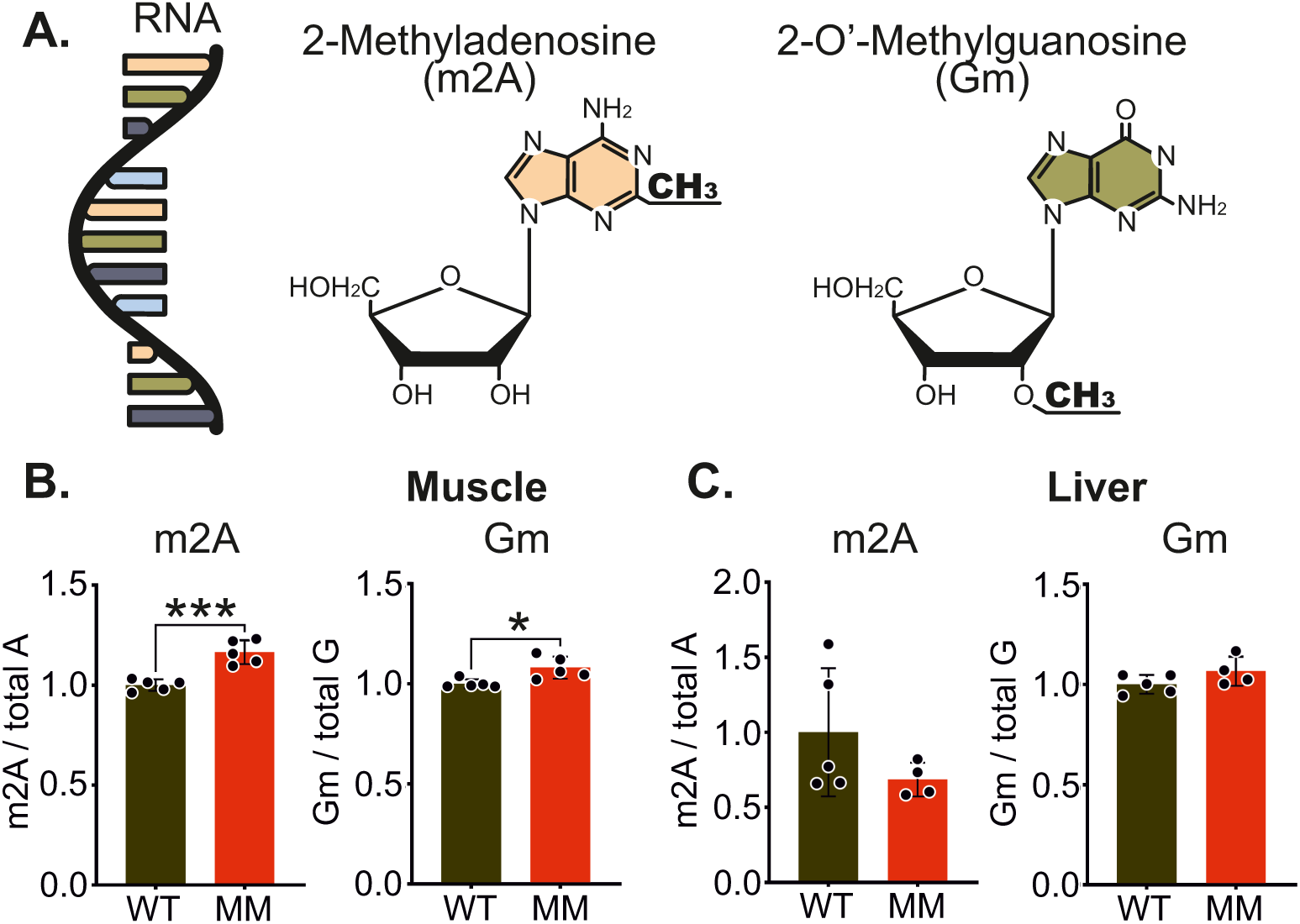
– Changes in RNA-specific methylation marks in MM muscles and livers. **(A)** Chemical structures of RNA nucleoside methylation marks. **(B-C)** Changes in methylated nucleoside levels, shown as the ratio of each methylated form to its corresponding total nucleoside level in (B) skeletal muscle and (C) liver (n = 5/group). Abbreviations: WT, wild type; MM, mitochondrial myopathy. Data information: Data are presented as mean ± SD. *P≤0.05, **P≤0.01, ***P≤0.001, ****P≤0.0001 (Student’s t-test)

Among the detected marks, m2A is the most understudied and has not been previously reported in mammalian RNAs. By contrast, Gm is one of the more abundant 2’-O-methylation marks. In rRNA and snRNA, site-specific Gm residues rigidify rRNA structure, promote proper ribosome assembly and translational fidelity, and ensure accurate spliceosome function (Natchiar et al, 2017; reviewed in Höfler & Carlomagno, 2020; Watkins & Bohnsack, 2012). In tRNAs, Gm modification in the anticodon loop enforces correct codon-anticodon pairing (Guy et al, 2015). Increase in these marks in MM may potentially serve as an additional mechanism regulating the changing translation processes required for adaptation to the dynamic metabolic changes.

Although 2’-O-methylation also occurs on mRNA (Li et al, 2024), our analysis used total RNA without fractionation. Consequently, any mRNA-derived signal is likely to be difficult to detect, and the profile predominantly reflects rRNA and tRNA. m2A, on the other hand, was long considered unique to bacteria but has recently been detected in plant chloroplast rRNA and tRNA (Duan et al, 2024). Its occurrence in mammalian cells remains largely uncharacterized, thus confirming the identity of this peak by LC-MS/MS will be an important next step. Collectively, these observations raise the possibility that altered RNA methylation contributes to post-transcriptional control of gene expression in MM, yet this hypothesis remains to be tested directly.

All in all, this study constructs a detailed profile of methylation changes specific for the skeletal muscle tissue in the context of muscle-specific disease with liver serving as a control tissue less affected by the disorder. Our findings show that mtDNA replication defect in the skeletal muscle causes robust metabolic changes of the methyl cycle with decreased PC and creatine synthesis and disease-specific chromatin modifications.

A large part of these methyl groups is deposited in the form of H3K9me3 histone mark and to a lesser extent in RNA. Still, ISRmt gene expression was not regulated by this mark.

Post-transcriptional RNA modifications are emerging regulators of gene expression. We find that 2’-O-methylation at guanosine (Gm) is increased in MM muscle. These changes could potentially modulate translation via rRNA and tRNA and influence pre-mRNA splicing via snRNA.

Ultimately, our studies highlight tissue-specific metabolism in MM. Understanding such specifics allows to develop disease-tailored therapies targeted to the organ of interest.

## Supporting information

Expanded view

## Figure legends

**Figure EV1 (A)** mRNA expression of MAT1a, PEMT, and AHCY enzymes in liver (n=8-9/group). **(B)** mRNA expression of MAT2a and AHCY enzymes in skeletal muscle (n=9/group). Abbreviations: MAT2a/1a, Methionine adenosyltransferase 2a/1a; AHCY, Adenosylhomocysteinase; PEMT, Phosphatidylethanolamine N-methyltransferase; WT, wild type; MM, mitochondrial myopathy.

Data information: Data are presented as 25^th^ and 75^th^ percentiles with whiskers spanning to min and max points. *P≤0.05, **P≤0.01, ***P≤0.001, ****P≤0.0001 (Student’s t-test)

**Figure EV2 (A-B)** Gene ontology analysis for (A) Biological process and (B) Cellular Component of detected ChIP-Seq peaks in skeletal muscle. **(C)** Metagene profile of H3K9me3 occupancy at representative differential H3K9me3 regions annotated to genes from Fig. 3D. **(D)** Chromatin state distribution for up- and down-regulated peak regions.

Abbreviations: mHet – heterochromatin; mQuies – quiescent; mGapArtf - assembly gaps and artifacts; mZNF - Zinc finger genes; mEnhWk - weak enhancers; mEnhA - active enhancers; mTxWk - weak transcription; mOpenC - open chromatin; mReprPC - polycomb repressed; mTxEx - transcription and exon.

**Figure EV3 (A-B)** Levels RNA methylation marks, shown as the ratio of each methylated form to its corresponding total nucleoside level in (A) skeletal muscle and (B) liver. Abbreviations: m6A, N6-methyladenosine; (m6)2A, N6,N6-dimethyladenosine; m1G, N1-methylguanosine; m2G, N2-methylguanosine; Cm, 2′-O-methylcytidine; m3C, N3-methylcytidine; m5C, 5-methylcytidine; m5U, 5-methyluridine; WT, wild type; MM, mitochondrial myopathy.

Data information: Data are presented as mean ± SD. *P≤0.05, **P≤0.01, ***P≤0.001, ****P≤0.0001 (Student’s t-test)

## Materials and Methods

### Ethical approval

All the experimental procedures and handling of mice were performed according to the national and international guidelines. License number ESAVI/11682/04.10.07/2017 approved by the Finnish Committee of Experimental Animal research.

### Animal models

This study utilized wild type and Deletor animals that were littermates of the same congenic strain – C57BL/6. Wild types expressed normal version of mitochondrial Twinkle helicase gene (Twnk). Deletors overexpressed Twnk transgene containing a patient dominant point mutation c.1077C>G, that leads to an in-frame duplication of 353-365. Male animals show more pronounced myopathy phenotype and hence were used in the study. Age of 22 month was chosen, when the disease is presents full phenotypical changes and is in chronic stage.

Animal genotype abbreviations: WT – wild type; DEL – deletor.

Housing of animals was at +25 °C, controlled light/dark cycle of 12 hours, food availability ad libitum.

### Tissue collection

To control for the similar food uptake, mice were fasted overnight (food removed the previous day at 7 pm) and refed at 6 am prior to termination. Diet – standard Chow. Animals were sacrificed with CO2 and neck dislocation. Collected organs were flash frozen in liquid nitrogen and stored at –80 °C until use.

### Targeted metabolomics

Detailed description of the targeted metabolomics procedure and reagents can be found in Nikkanen et al (2016). Briefly, prior to measurements, the instrument was calibrated for the 100-metabolite panel: a 10-point water-based serial dilution was prepared in a 96-well plate from a stock mix (automated pipetting Hamilton MICROLAB® STAR line). Metabolite extraction was performed from frozen tissue (muscle and liver, 15–35 mg), which was mixed with 20 µL labelled internal-standard mix and 1:30 (sample:solvent) extraction buffer and extracted in two steps using bead homogenization. In step 1, 15 vol of ice-cold 100% ACN with 1% Formic acid (FA) was added to the sample and homogenized at 5500 rpm for 20 sec 3 times with 30 sec pauses. The debris and supernatant were separated by centrifugation at 5000 rpm for 10 min at +4 °C; supernatant collected into a separate tube. In step 2, 15 vol of 80:20% ACN:H₂O with 1% FA was added to the pellet and the homogenization was repeated. The extracts of both steps were pooled and centrifuged at 5,000 rpm for 15 min at +4 °C; collect clarified extract.

Metabolomics data were processed and analyzed with the web-based MetaboAnalyst 3.0 platform (Xia et al, 2015). Analyses used non-transformed values with autoscaling (mean-centering and division by each variable’s standard deviation).

### Quantitative mass spectrometry of glycerophospholipids

The methodology for the mass-spectrometric analysis of lipids was described previously (Özbalci et al, 2013; Tatsuta, 2017) and the dataset used has been published (Jackson et al, 2025).

Briefly, lipids were extracted from a mitochondria-enriched fraction using a modified Bligh–Dyer protocol in the presence of internal standards for the major phospholipid classes including PC 17:0/20:4 and PE 17:0/20:4. Extracts were analyzed on a QTRAP 6500 triple-quadrupole mass spectrometer (SCIEX) equipped with a TriVersa NanoMate nano-infusion ESI source with type-A chip (Advion). Mass spectra were processed in LipidView v1.2 (SCIEX) for lipid identification and quantification. Reported amounts (pmol) were corrected for response differences between internal standards and endogenous species.

### RNA isolation and RT-qPCR

RNA extraction was done using miRNeasy kit with DNase treatment according to the manufacturer’s instructions. For RT-qPCR, the quantity and quality of the RNA were evaluated with NanoDrop spectrophotometer. cDNA synthesis was done Maxima First Strand cDNA Synthesis Kit according to the manufacturer’s instructions. The RT reaction was done using SensiFAST™ SYBR No-ROX Kit with the reaction protocol as follows:

Step 1: 2 min at 95 °C polymerase activation
Step 2: 5 sec at 95 °C denaturation
Step 3: 30 sec at 60 °C annealing and extension
Repeated steps 2-3: total of 40 cycles
Step 4: 65 - 95 °C increase with 0.5 °C increments for melt curve

### ChIP-Sequencing

The whole process of ChIP-Sequencing required a 4-day long protocol. It is essential to keep samples and buffers in cold (+4 or –80 °C) unless specified otherwise. Below the steps are characterised as per one sample.

### PBS/BSA

#### DAY 1 – Antibody coupling to magnetic beads

Collection of QF muscles from mice was done as described above. Tissues were flash frozen in liquid nitrogen and stored at -80°C until use in the experiments.

Buffers were freshly prepared and stored at +4°C.

50 µL magnetic beads were resuspended and washed 3 times in 1 mL PBS/BSA using a magnetic rack. The beads were resuspended again in 1 mL PBS/BSA with 5 µg/mL antibodies – IgG or H3K9me3 – and gently mixed on a rotator platform overnight at +4 °C.

#### DAY 2 – Tissue preparation, cross-linking and immunoprecipitation

Frozen muscles were homogenized with mortar and pestle on dry ice. Tissue powder was resuspended in 2 mL of room temperature (RT) PBS with protease inhibitor cocktail (PIC). The suspension was then transferred to 15 mL conical tubes and volume brought up to 10 mL of PBS-PIC. The cells were crosslinked in 1% formaldehyde for 15 min on a rocker at RT. The crosslinking reaction was blocked with 0.125 M glycine. The sample was washed 3 times in 10 mL cold PBS-PIC for 5 min at 500 g at +4 °C.

After the final wash, the pellet was resuspended in 500 µL PB buffer, transferred into fresh tubes with Precellys Mixed Beads (Cat.no. P000918-LYSK0-A, Precellys) and further homogenized at 5500 rpm for 15 seconds with Precellys 24 bead homogenizer. The homogenate was transferred into new tubes without beads for centrifugation at 6300 g for 10 min at +4 °C. The supernatant was discarded, and the pellet was resuspended in 500 µL RIPA buffer with PIC. The samples were stored on ice until the sonication step.

Sonication was done using (Otonkoski) sonicator for 40 cycles with the following settings: output – 9.5 W; sonication ON – 20 sec, OFF – 60 sec. Sonicated samples were centrifuged at 13200 rpm for 15 min at +4 °C, and the supernatant was transferred into fresh tubes.

IP reactions were set: 100 µL of sonicated chromatin was mixed with 900 µL of RIPA-PIC, of which 50 µL was removed as an input control. The antibody-coupled beads were washed 3 times with 1 mL PBS-BSA. Washed beads were resuspended in 100 µL PBS-BSA and added to the IP reaction. The IP mix was incubated on the rotating mixer overnight at +4 °C.

#### DAY 3 – Washes and reverse cross-linking

IP reactions were washed 5 times with LiCl wash buffer for 3 min at +4 °C followed by 2 washes in TE buffer for 1 min at +4 °C. Washed beads were resuspended in 200 µL of elution buffer and heated at 65 °C for 1 hr with constant shaking at 600 rpm. After this, the magnetic beads were separated, and a clear supernatant was transferred to a fresh tube. Proteinase K was added to the IP samples at the final concentration of 200 µg/mL and the samples were incubated again at +65 °C overnight.

#### DAY 4 – DNA purification

IP samples were diluted in equal volume of TE buffer with 200 µg/mL RNAase A (Cat.no. EN0531, Thermo Fisher Scientific) and incubated at +37 °C for 1 hr. DNA from both the IP reaction and input was isolated with standard phenol-chloroform protocol and eluted in 100 µL elution buffer. The samples were stored at –80 °C until being shipped for sequencing.

### ChIP-Sequencing and data analysis

Quality check of the samples, library preparation and sequencing were done at BGI Genomics. Sequencing with DNBSEQ technology, depth of 50 million, paired end, 100 gb reads.

In short, ChIP-seq data quality was assessed using fastqc v.0.11.8 (https://www.bioinformatics.babraham.ac.uk/projects/fastqc/). Single-end reads were aligned to the mm10 reference genome using Bowtie2 v.2.2.5 [bowtie2 --very-sensitive] (Langmead & Salzberg, 2012). Duplicate reads were identified using Picard v.2.26.3 (https://broadinstitute.github.io/picard/) [picard MarkDuplicates - REMOVE_DUPLICATES false -ASSUME_SORT_ORDER coordinate] and removed, along with low-quality reads (MAPQ <20), using samtools v.1.9 [samtools view –F 1024 –q 20] (Li et al, 2009). Reads were sorted using samtools (samtools sort). Peaks were called using Macs2 v.2.1.1.20160309 [macs2 callpeak -g mm --broad --keep-dup all] with IgG controls (Zhang et al, 2008; Feng et al, 2012). ENCODE blacklisted regions (accession: ENCFF547MET) (Amemiya et al, 2019) were removed from the broad peaks using bedtools v.2.30.0 (Quinlan & Hall, 2010). Normalized signal files were generated using deepTools v.3.1.3 [bamCoverage --normalizeUsing RPKM -bl ENCFF547MET.bed -- binSize 50] (Ramírez et al, 2014, 2016).

Peak files were further analyzed in R (v4.2.2 and above) using a set of packages. ChIPpeakAnno (v3.32.0; Zhu et al, 2010) was used to annotate peaks to the nearby genes using TxDb.Mmusculus.UCSC.mm10.knownGene annotation, to find overlapping peaks between replicates, to define genomic element distribution of peaks, and to run gene ontology analysis on the full WT and DEL peaksets. Differential binding analysis was done with DiffBind (v3.12.0; Stark & Brown, 2011), using a consensus peakset with significance determined at FDR < 0.05. Sites in the ENCODE blacklisted regions were removed. Differential regions were annotated to their nearest genes and genomic features with ChIPseeker (v1.34.1; Yu et al, 2015) and to the chromatin state using ChromHMM annotation (Ernst & Kellis, 2012). GO enrichment of these regions was analyzed with clusterProfiler (v4.10.1; Yu et al, 2012; Wu et al, 2021). Additionally, graphics were created using ggplot2 package (v3.5.2; Wickham, 2016). Metagene profiles were generated using ggplot2 showing mean +/- CI (t-distribution, n=3) across replicates, binned either around the called peak (+/-0.5 x peak width) or across the full gene body (20 bins/gene).

### Protein extraction and immunoblotting

An aliquot of nuclei in RIPA buffer was preserved from the nuclear isolation stage of ChIP-sequencing protocol. The sample was briefly sonicated to break down the membrane and the protein amount was measured with Bradford assay (Cat.no. 5000006, Bio-Rad). The protein was resuspended in Laemmli buffer and heated at 95 °C for 15 min.

6 µL of protein sample was then loaded and separated on 4-20% Stain-Free SDS gel (Cat.no. 5678094, Bio-Rad) and transferred onto PVDF membrane with semi-dry method (Trans-Blot Turbo, Bio-Rad). The membranes were blocked in 5% milk in TBST (Tris Buffered Saline with 0.1% Tween20) for 1 hr at RT, after which they were incubated with primary antibodies overnight at +4 °C. Concentrations of primary antibodies were applied according to manufacturer’s instructions. The membranes were washed 3 times for 10 min in TBST the next day. Secondary HRP-conjugated antibodies were diluted 1 to 10000 in 5% milk in TBST and incubated with the membranes for 1 hr at RT. Prior to imaging, the membranes were washed again in TBST as before. The signal was visualised with Clarity Western ECL Substrate (Cat.no. 1705061, Bio-Rad) and ChemiDoc™ XRS+ System (Biorad). Normalization and quantification of signal intensities were done in ImageLab 6.1.0 (Bio-Rad).

### Detection of methylated RNA

First, total RNA was extracted from QF and liver tissue samples. Tissues (40-60mg) were homogenized in 1 mL Trizol (Cat.no. 15596018, Ambion) with bead homogeniser. Precellys Lysin kit with beads for hard tissue (KT03961-1-002.2, Fisher Scientific) were used for muscle and soft tissue (KT03961-1-009.2, Fisher Scientific) – for liver.

Homogenization settings: 5000 rpm x 2 cycles for 30 sec ach and 60 seconds rest for muscle; 5500 rpm x 1 cycle for 15 sec for liver. After homogenization, the samples were incubated at RT for 5 min, after which the homogenate was transferred into clean tubes with 200 µL chloroform, shook for 15 sec, incubated at RT for 2-3 min and centrifuged at 12000 g, 15 min, +4 °C. The upper aqueous phase was transferred and mixed with equal amount of isopropanol and incubated at RT for 10 min. The mix was centrifuges again at 12000 g, 10 min, +4 °C to precipitate an RNA pellet. The pellet was washed first with 75% ethanol and centrifugation at 7500 g, 5 min, +4 °C after which the ethanol was removed, and the pellet was left to dry under the clean hood for 10 min. RNA pellet was resuspended in 180 µL DEPC-treated water for further steps.

The freshly isolated total RNA then underwent DNase treatment and cleanup. For this, it was first incubated at +55 °C for 10 min. DNase reaction was setup by adding DNase buffer to 1x concentration and 10 units of DNase I enzyme (Cat.no. 79254, Qiagen) per reaction. The mix was incubated for 1 hr at +37 °C. At the end of the reaction, RNA was purified by mixing it with 1:1 volume of phenol:chloroform:isoamyl alcohol and centrifugation at maximum speed, 5 min, +4 °C. The upper phase was transferred into a fresh tube, mixed with 1 volume of chloroform and centrifuged as in the previous step. The upper phase was collected separately again, and RNA was precipitated in 1/10 volume of 3M NaAc, 2.5 volumes of cold 95% EtOH and incubation at –20 °C overnight. RNA is recovered by centrifugation at 11000 rpm, 10 min, +4 °C. The pellet was washed with 160 µL of 70% EtOH and centrifuged again at 11000 rpm, 10 min, +4 °C. The pellet was air dried in a clean hood for 5-10 min at RT, after which it was resuspended in the appropriate amount of DEPC-treated water and the concentration was measured with Qubit assay (Cat.no. Q10210, Invitrogen).

At the last stage of sample preparation, RNA was lysed into mononucleotides with P1 nuclease as follows. 40µg of purified RNA was brought to 82 µL volume with DEPC-treated water and heated at 95 °C for 4 min. The samples were allowed to cool down to RT. The enzyme reaction was set up by mixing the RNA with 760 µM ZnSO4 and 3-4 Units of P1 enzyme (Cat.no. N8630-1VL, Sigma Aldrich) with incubation at +37 °C overnight. The nucleosides were dephosphorylated in 1x FastAP buffer and 1 Unit of FastAP enzyme (Cat.no. EF0651, Thermo Fisher) at +37 °C for 3 hours.

The resulting nucleosides were analysed by reversed-phase HPLC using a modification of the method of Gehrke and Kuo (1989), essentially as described by Siibak and Remme (2010) and O’Connor et al (2018). Nucleosides were separated on a Supelcosil LC-18-S reversed-phase column using a Shimadzu Prominence HPLC system. Mobile phase A consisted of 10 mM NH₄H₂PO₄ containing 2.5% methanol (pH 5.3), mobile phase B of 10 mM NH₄H₂PO₄ containing 20% methanol (pH 5.1), and mobile phase C of 10 mM NH₄H₂PO₄ containing 35% acetonitrile (pH 4.9). The column was maintained at 30 °C and operated at a flow rate of 1.0 mL min⁻¹. Absorbance was monitored at 260 and 280 nm. Modified nucleosides were assigned on the basis of their relative retention times according to Gehrke and Kuo (1989).

## Data availability

RNA methylation data has been deposited to Zenodo doi: 10.5281/zenodo.22050816.

## Acknowledgements

We thank Gulayse Ince-Dunn and Liliya Euro, Margus Leppik, Joni Nikkanen and Takashi Tatsuta for technical contributions; Saara Forsström for comments and feedback on the early drafts of the manuscript; Metabolomics Unit, Institute for Molecular Medicine Finland (FIMM) for metabolomics data; BGI Genomics for ChIP-Sequencing. We also thank Finnish Cultural Foundation (to Anastasiia Marmyleva # 00210706) and Magnus Ehrnrooth Foundation (to Anastasiia Marmyleva) for the financial support of this work.

## Author information

### Contributions

**Anastasiia Marmyleva:** Conceptualization; Data curation; Formal analysis; Funding acquisition; Investigation; Methodology; Project administration; Validation; Visualization; Writing—original draft; Writing—review and editing. **Ville Tiusanen:** Data curation; Formal analysis; Validation; Writing—review and editing. **Priit Joers:** Investigation; Methodology; Writing—review and editing. **Biswajyoti Sahu:** Methodology; Validation; Writing—review and editing. **Anu Suomalainen:** Conceptualization; Funding acquisition; Supervision; Writing—review and editing.

### Corresponding author

Correspondence to Anu Suomalainen.

### Ethics declaration

The authors declare no competing interests.

## References

Amemiya HM, Kundaje A, Boyle AP (2019) The ENCODE blacklist: identification of problematic regions of the genome. Sci Rep 9: 9354

Bannister AJ, Kouzarides T (2011) Regulation of chromatin by histone modifications. Cell Res 21: 381–395

Bao XR, Ong SE, Goldberger O, Peng J, Sharma R, Thompson DA, Vafai SB, Cox AG, Marutani E, Ichinose F et al (2016) Mitochondrial dysfunction remodels one-carbon metabolism in human cells. Elife 5: e10575

Buzkova J, Nikkanen J, Ahola S, Hakonen AH, Sevastianova K, Hovinen T, Yki-Järvinen H, Pietiläinen KH, Lönnqvist T, Velagapudi V et al (2018) Metabolomes of mitochondrial diseases and inclusion body myositis patients: treatment targets and biomarkers. EMBO Mol Med 10: e9091

Duan HC, Zhang C, Song P, Yang J, Wang Y, Jia G (2024) C2-methyladenosine in tRNA promotes protein translation by facilitating the decoding of tandem m2A-tRNA-dependent codons. Nat Commun 15: 1025

Ernst J, Kellis M (2012) ChromHMM: automating chromatin-state discovery and characterization. Nat Methods 9: 215–216

Feng J, Liu T, Qin B, Zhang Y, Liu XS (2012) Identifying ChIP-seq enrichment using MACS. Nat Protoc 7: 1728–1740

Forsström S, Jackson CB, Carroll CJ, Kuronen M, Pirinen E, Pradhan S, Marmyleva A, Auranen M, Kleine IM, Khan NA et al (2019) Fibroblast growth factor 21 drives dynamics of local and systemic stress responses in mitochondrial myopathy with mtDNA deletions. Cell Metab 30: 1040–1054

Gehrke CW, Kuo KC (1989) Ribonucleoside analysis by reversed-phase high-performance liquid chromatography. J Chromatogr 471: 3–36

Guy MP, Shaw M, Weiner CL, Hobson L, Stark Z, Rose K, Kalscheuer VM, Gecz J, Phizicky EM (2015) Defects in tRNA anticodon loop 2’-O-methylation are implicated in nonsyndromic X-linked intellectual disability due to mutations in FTSJ1. Hum Mutat 36: 1176–1187

Hada M, Miura H, Tanigawa A, Matoba S, Inoue K, Ogonuki N, Hirose M, Watanabe N, Nakato R, Fujiki K et al (2022) Highly rigid H3.1/H3.2-H3K9me3 domains set a barrier for cell fate reprogramming in trophoblast stem cells. Genes Dev 36: 84–102

Höfler S, Carlomagno T (2020) Structural and functional roles of 2’-O-ribose methylations and their enzymatic machinery across multiple classes of RNAs. Curr Opin Struct Biol 65: 42–50

Jackson CB, Marmyleva A, Monteuuis G, Awadhpersad R, Mito T, Zamboni N, Tatsuta T, Vincent AE, Wang L, Khan NA et al (2025) De novo serine biosynthesis is protective in mitochondrial disease. Cell Rep 44: 115710

Karimian A, Vogelauer M, Kurdistani SK (2023) Metabolic sinkholes: histones as methyl repositories. PLoS Biol 21: e3002371

Khan NA, Nikkanen J, Yatsuga S, Jackson C, Wang L, Pradhan S, Kivelä R, Pessia A, Velagapudi V, Suomalainen A (2017) mTORC1 regulates mitochondrial integrated stress response and mitochondrial myopathy progression. Cell Metab 26: 419–428

Kühl I, Miranda M, Atanassov I, Kuznetsova I, Hinze Y, Mourier A, Filipovska A, Larsson NG (2017) Transcriptomic and proteomic landscape of mitochondrial dysfunction reveals secondary coenzyme Q deficiency in mammals. Elife 6: e30952

Lachner M, O’Carroll D, Rea S, Mechtler K, Jenuwein T (2001) Methylation of histone H3 lysine 9 creates a binding site for HP1 proteins. Nature 410: 116–120

Langmead B, Salzberg SL (2012) Fast gapped-read alignment with Bowtie 2. Nat Methods 9: 357–359

Lee MY, Lee J, Hyeon SJ, Cho H, Hwang YJ, Shin JY, McKee AC, Kowall NW, Kim JI, Stein TD et al (2020) Epigenome signatures landscaped by histone H3K9me3 are associated with the synaptic dysfunction in Alzheimer’s disease. Aging Cell 19: e13153

Li H, Handsaker B, Wysoker A, Fennell T, Ruan J, Homer N, Marth G, Abecasis G, Durbin R, 1000 Genome Project Data Processing Subgroup (2009) The Sequence Alignment/Map format and SAMtools. Bioinformatics 25: 2078–2079

Li Y, Yi Y, Gao X, Wang X, Zhao D, Wang R, Zhang LS, Gao B, Zhang Y, Zhang L et al (2024) 2’-O-methylation at internal sites on mRNA promotes mRNA stability. Mol Cell 84: 2320–2336

Methot SP, Padeken J, Brancati G, Zeller P, Delaney CE, Gaidatzis D, Kohler H, van Oudenaarden A, Großhans H, Gasser SM (2021) H3K9me selectively blocks transcription factor activity and ensures differentiated tissue integrity. Nat Cell Biol 23: 1163–1175

Mudd SH, Brosnan JT, Brosnan ME, Jacobs RL, Stabler SP, Allen RH, Vance DE, Wagner C (2007) Methyl balance and transmethylation fluxes in humans. Am J Clin Nutr 85: 19–25

Natchiar SK, Myasnikov AG, Kratzat H, Hazemann I, Klaholz BP (2017) Visualization of chemical modifications in the human 80S ribosome structure. Nature 551: 472–477

Nikkanen J, Forsström S, Euro L, Paetau I, Kohnz RA, Wang L, Chilov D, Viinamäki J, Roivainen A, Marjamäki P et al (2016) Mitochondrial DNA replication defects disturb cellular dNTP pools and remodel one-carbon metabolism. Cell Metab 23: 635–648

O’Connor M, Leppik M, Remme J (2018) Pseudouridine-free Escherichia coli ribosomes. J Bacteriol 200: e00540–17

Özbalci C, Sachsenheimer T, Brügger B (2013) Quantitative analysis of cellular lipids by nano-electrospray ionization mass spectrometry. Methods Mol Biol 1033: 3–20

Peters AH, Kubicek S, Mechtler K, O’Sullivan RJ, Derijck AA, Perez-Burgos L, Kohlmaier A, Opravil S, Tachibana M, Shinkai Y et al (2003) Partitioning and plasticity of repressive histone methylation states in mammalian chromatin. Mol Cell 12: 1577–1589

Pirinen E, Auranen M, Khan NA, Brilhante V, Urho N, Pessia A, Hakkarainen A, Kuula J, Heinonen U, Schmidt MS et al (2020) Niacin cures systemic NAD+ deficiency and improves muscle performance in adult-onset mitochondrial myopathy. Cell Metab 31: 1078–1090

Quinlan AR, Hall IM (2010) BEDTools: a flexible suite of utilities for comparing genomic features. Bioinformatics 26: 841–842

Ramírez F, Dündar F, Diehl S, Grüning BA, Manke T (2014) deepTools: a flexible platform for exploring deep-sequencing data. Nucleic Acids Res 42: W187–W191

Ramírez F, Ryan DP, Grüning B, Bhardwaj V, Kilpert F, Richter AS, Heyne S, Dündar F, Manke T (2016) deepTools2: a next generation web server for deep-sequencing data analysis. Nucleic Acids Res 44: W160–W165

Schlesinger S, Sonntag SR, Lieb W, Maas R (2016) Asymmetric and symmetric dimethylarginine as risk markers for total mortality and cardiovascular outcomes: a systematic review and meta-analysis of prospective studies. PLoS One 11: e0165811

Siibak T, Remme J (2010) Subribosomal particle analysis reveals the stages of bacterial ribosome assembly at which rRNA nucleotides are modified. RNA 16: 2023–2032

Soufi A, Donahue G, Zaret KS (2012) Facilitators and impediments of the pluripotency reprogramming factors’ initial engagement with the genome. Cell 151: 994–1004

Stark R, Brown G (2011) DiffBind: differential binding analysis of ChIP-Seq peak data. Bioconductor package version 3.12.0. https://bioconductor.org/packages/3.18/bioc/vignettes/DiffBind/inst/doc/DiffBind.pdf

Suomalainen A, Battersby BJ (2018) Mitochondrial diseases: the contribution of organelle stress responses to pathology. Nat Rev Mol Cell Biol 19: 77–92

Suomalainen A, Nunnari J (2024) Mitochondria at the crossroads of health and disease. Cell 187: 2601–2627

Tatarakis A, Saini H, Yu J, Feng W, Pinzon-Arteaga CA, Moazed D (2025) Requirements for establishment and epigenetic stability of mammalian heterochromatin. Mol Cell 85: 3388–3406

Tatsuta T (2017) Quantitative analysis of glycerophospholipids in mitochondria by mass spectrometry. Methods Mol Biol 1567: 79–103

Tyynismaa H, Mjosund KP, Wanrooij S, Lappalainen I, Ylikallio E, Jalanko A, Spelbrink JN, Paetau A, Suomalainen A (2005) Mutant mitochondrial helicase Twinkle causes multiple mtDNA deletions and a late-onset mitochondrial disease in mice. Proc Natl Acad Sci USA 102: 17687–17692

Tyynismaa H, Carroll CJ, Raimundo N, Ahola-Erkkilä S, Wenz T, Ruhanen H, Guse K, Hemminki A, Peltola-Mjøsund KE, Tulkki V et al (2010) Mitochondrial myopathy induces a starvation-like response. Hum Mol Genet 19: 3948–3958

Verschure PJ, van der Kraan I, de Leeuw W, van der Vlag J, Carpenter AE, Belmont AS, van Driel R (2005) In vivo HP1 targeting causes large-scale chromatin condensation and enhanced histone lysine methylation. Mol Cell Biol 25: 4552–4564

Watkins NJ, Bohnsack MT (2012) The box C/D and H/ACA snoRNPs: key players in the modification, processing and the dynamic folding of ribosomal RNA. Wiley Interdiscip Rev RNA 3: 397–414

Wickham H (2016) ggplot2: elegant graphics for data analysis, 2nd edn. Cham: Springer International Publishing

Wu T, Hu E, Xu S, Chen M, Guo P, Dai Z, Feng T, Zhou L, Tang W, Zhan L et al (2021) clusterProfiler 4.0: a universal enrichment tool for interpreting omics data. Innovation (Camb) 2: 100141

Xia J, Sinelnikov IV, Han B, Wishart DS (2015) MetaboAnalyst 3.0 - making metabolomics more meaningful. Nucleic Acids Res 43: W251–W257

Yu G, Wang LG, Han Y, He QY (2012) clusterProfiler: an R package for comparing biological themes among gene clusters. OMICS 16: 284–287

Yu G, Wang LG, He QY (2015) ChIPseeker: an R/Bioconductor package for ChIP peak annotation, comparison and visualization. Bioinformatics 31: 2382–2383

Zhang Y, Liu T, Meyer CA, Eeckhoute J, Johnson DS, Bernstein BE, Nusbaum C, Myers RM, Brown M, Li W et al (2008) Model-based analysis of ChIP-Seq (MACS). Genome Biol 9: R137

Zhao E, Ding J, Xia Y, Liu M, Ye B, Choi JH, Yan C, Dong Z, Huang S, Zha Y et al (2016) KDM4C and ATF4 cooperate in transcriptional control of amino acid metabolism. Cell Rep 14: 506–519

Zhu LJ, Gazin C, Lawson ND, Pagès H, Lin SM, Lapointe DS, Green MR (2010) ChIPpeakAnno: a Bioconductor package to annotate ChIP-seq and ChIP-chip data. BMC Bioinformatics 11: 237

