## Supplementary figures and images for "Mitochondrial dysfunction reshapes methyl-group allocation in skeletal muscle"

### Expanded view

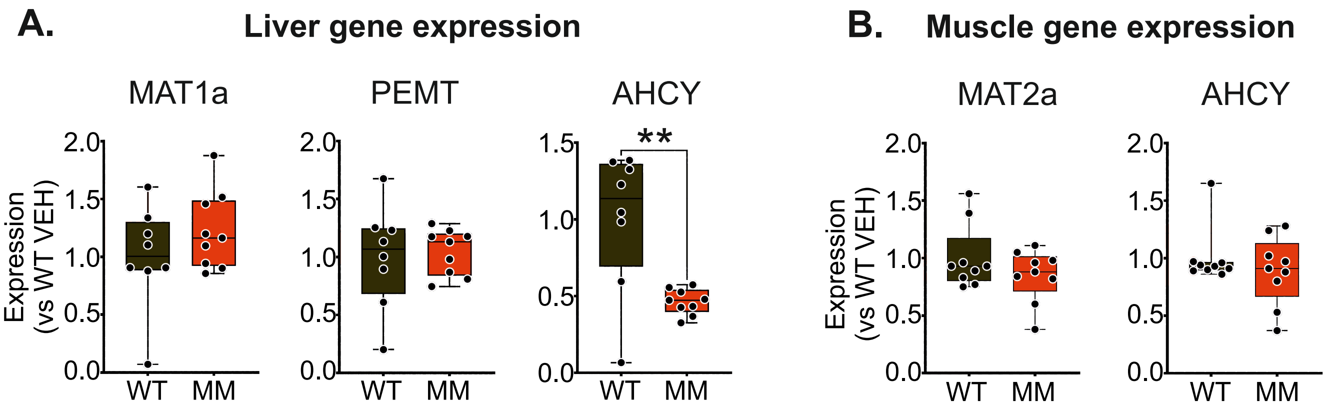

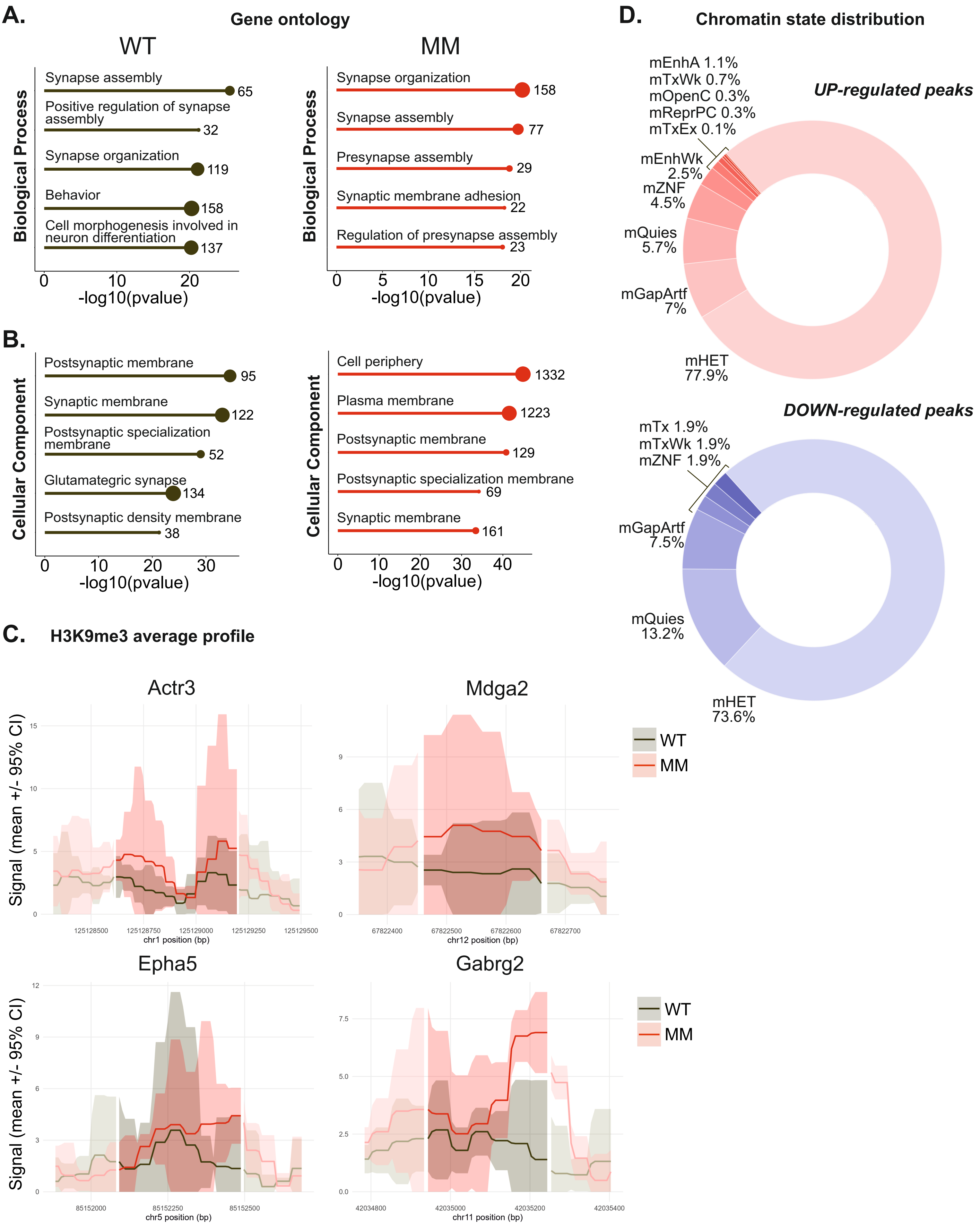

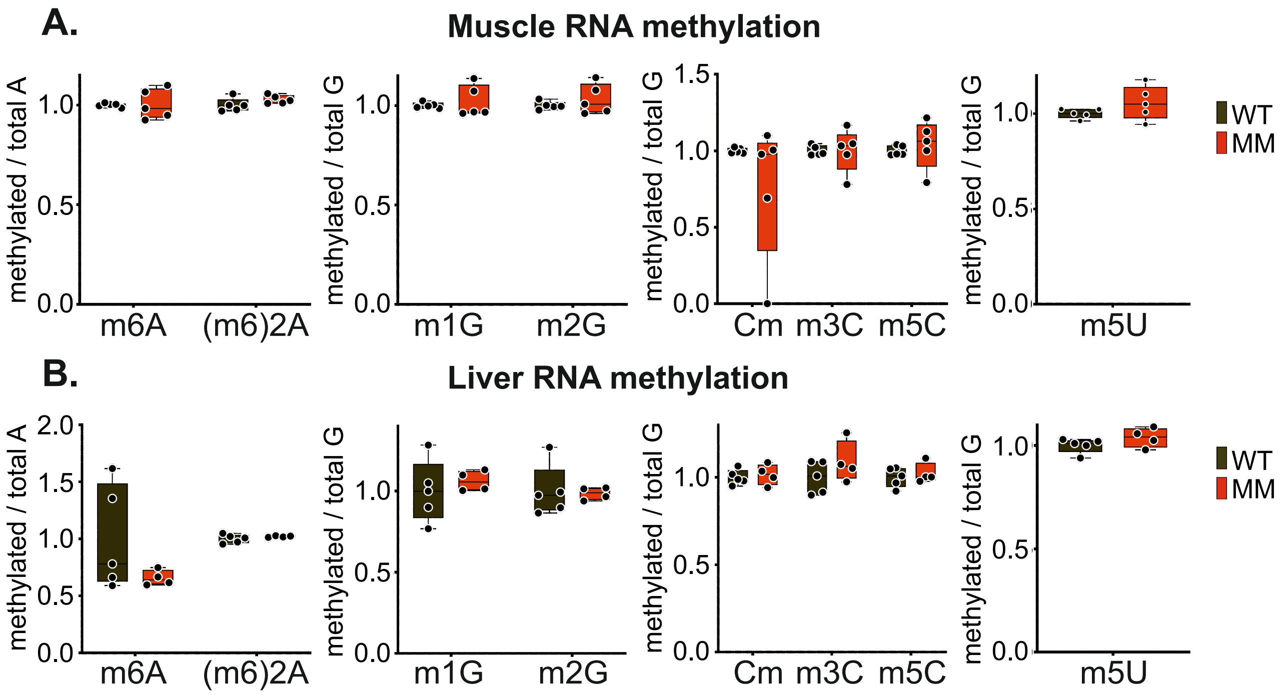
